# A Novel Cyanine-Based Fluorescent Dye for Targeted Mitochondrial Imaging in Neurotoxic Conditions and *In Vivo* Brain Studies

**DOI:** 10.1101/2024.10.24.619902

**Authors:** Xin Yan, Xinqian Chen, Shanshan Hou, Xiaoxu Li, Jingyue Ju, Zhiying Shan, Lanrong Bi

## Abstract

Mitochondrial dysfunction is a key feature of neurodegenerative diseases, often preceding symptoms and influencing disease progression. However, real-time *in vivo* imaging of mitochondria in the brain is limited by existing dyes like MitoTrackers, which struggle with poor tissue penetration, phototoxicity, and inability to cross the blood-brain barrier (BBB). This study introduces Cy5-PEG4, a novel mitochondrial-targeting dye that overcomes these limitations, enabling high-resolution, non-invasive imaging of mitochondrial dynamics. Cy5-PEG4 effectively labels mitochondria in primary neuronal cells exposed to the SARS-CoV-2 RNYIAQVD peptide, revealing dose-dependent alterations in mitochondrial function that may contribute to COVID-19-related neurodegeneration. Importantly, Cy5-PEG4 crosses the BBB without causing neuroinflammation or toxicity, making it a safe tool for *in vivo* brain imaging and detailed studies of mitochondrial responses. In 3D cultured cells, Cy5-PEG4 captures dynamic changes in mitochondrial distribution and morphology as cell structures mature, highlighting its potential in neurobiological research, diagnostics, and therapeutic development. These findings support Cy5-PEG4 as a powerful tool for studying disease progression, identifying early biomarkers, and evaluating therapeutic strategies in neurodegenerative disorders and COVID-19.

## Introduction

Mitochondrial dysfunction is a key feature of many neurodegenerative diseases, such as Alzheimer’s, Parkinson’s, and Huntington’s, where changes in mitochondrial dynamics and morphology often precede clinical symptoms [1]. Imaging mitochondria within the living brain allows researchers to observe these organelles in their natural environment, leading to a more accurate understanding of their behavior, dynamics, and role in disease progression [2]. This capability is vital for monitoring mitochondrial changes, such as fission and fusion, which reflect cellular health and metabolic state. Traditional methods, relying on *in vitro* studies or post-mortem tissue analysis, cannot capture the dynamic nature of mitochondrial behavior during disease development. *In vivo* imaging with novel fluorescent dyes may bridge this gap, providing real-time insights into mitochondrial responses to pathological conditions, therapeutic interventions, and environmental changes.

MitoTrackers are extensively used for fluorescent staining to study mitochondrial dynamics and health in live and fixed cells. However, their application in brain cell imaging, especially *in vivo*, is significantly challenged by several factors [3]. Their optical properties in the visible light spectrum led to limited tissue penetration, resulting in poor imaging quality in deeper brain tissues. The blood-brain barrier (BBB) further restricts MitoTrackers from reaching brain cells, necessitating dye modifications or complex delivery methods that complicate experimental setups. MitoTrackers also present risks such as phototoxicity due to prolonged illumination during fluorescence imaging, which can impact mitochondrial function and cell viability in delicate brain tissues [3]. Additionally, brain tissues’ inherent autofluorescence can interfere with MitoTracker detection, masking mitochondrial signals, while photobleaching hampers the long-term observation of mitochondrial behavior. Since MitoTracker staining relies on mitochondrial membrane potential, it may not accurately represent mitochondrial quantity or health, particularly in pathological conditions where membrane potential is altered. Furthermore, the MitoTracker dyes themselves can disrupt mitochondrial function, potentially skewing experimental outcomes [4].

These limitations highlight the need for new dyes or imaging methodologies that can provide accurate, non-invasive examinations of mitochondria within brain cells during *in vivo* studies. Such advancements would offer more reliable metrics for the assessment of mitochondrial health and the support a deeper understanding of these organelles’ roles *in vivo*, underscoring the need for innovative solutions in neurological research. However, developing these novel fluorescent dyes presents significant challenges. The dense and complex environment of brain tissue dampens adequate dye penetration and precise imaging. Moreover, it adds complexity to design dyes that are non-toxic, stable, capable of crossing the BBB, and do not alter mitochondrial function upon binding. These dyes must also be compatible with existing imaging technologies and enable quantitative analysis that is essential for evaluating the bioenergetic health of neurons and understanding neurodegenerative disease progression.

This manuscript presents the synthesis and biological evaluation of a novel mitochondria-targeting fluorescent dye, Cy5-PEG4. We have successfully employed this dye to track changes in mitochondrial morphology and dynamics in primary neuronal cells exposed to the highly toxic RNYIAQVD peptide, a fragment associated with SARS-CoV-2 that is known to harm neural cells [5]. Furthermore, we investigated the capacity of Cy5-PEG4 to cross the BBB and assessed its potential for *in vivo* brain imaging.

## Material and Methods

The synthesis experiment was conducted using reagents and solvents sourced from commercial suppliers, which were used without further purification unless otherwise specified. Thin-Layer Chromatography (TLC) analyses were performed using silica gel TLC plates from Sigma-Aldrich, with a 0.25 µm coating thickness over glass support, denoted as plate number 1. Flash column chromatography utilized Alfa Aesar silica gel with a particle size range of 230–400 mesh. Melting points of compounds were determined using a MELTEMP apparatus without further correction. Nuclear magnetic resonance (NMR) spectroscopy was employed to obtain ^1^H and ^13^C NMR spectra, using a Varian UNITY INOVA instrument operating at frequencies of 400 MHz for ^1^H and 100 MHz for ^13^C, respectively. Chemical shifts (δ) were referenced to the residual peaks of the solvent dimethyl sulfoxide (DMSO)-d6, with ^1^H at δ = 2.50 ppm and ^13^C at δ = 39.52 ppm. NMR data were presented as follows: chemical shift, multiplicity (with s representing singlet, d for doublet, t for triplet, q for quartet, m for multiplet), coupling constants (J), and signal integration. High-resolution mass spectra (HR-MS) were acquired using a JEOL JMS HX 110A mass spectrometer. UV-visible (UV–vis) spectra were recorded at 37 °C on a PerkinElmer Lambda 35 UV/Vis Spectrometer, equipped with a PTP 1+1 Peltier Temperature Programmer accessory. Fluorescence spectra were measured with a Horiba Jobin Yvon Fluoromax-4 spectrofluorometer, setting the excitation wavelength at 360 nm and slit widths to 5 nm.

### Cell Culture

HEK293 cells, obtained from the American Type Culture Collection (ATCC), were grown in Dulbecco’s Modified Eagle’s Medium (DMEM) enriched with 10% Fetal Bovine Serum (FBS). The culture medium was refreshed every 2 to 3 days to promote optimal growth conditions.

Human umbilical vein endothelial cells (HUVEC), also purchased from ATCC, were cultured in DMEM. This medium was augmented with 5.5 mM glucose, 10% FBS, 100 U/ml penicillin, and 100 µg/ml streptomycin. Cultures were maintained at 37°C in a 5% CO_2_-humidified environment to ensure proper cell growth.

HeLa cells, sourced from ATCC, were cultured using RPMI-1640 medium supplemented with 10% FBS, 2 mM glutamine, and 100 µg/mL each of penicillin and streptomycin. These cells were kept in a 5% CO_2_ humidified atmosphere at a constant temperature of 37°C, specifically for imaging studies.

### Subcellular Localization Study of Cy5-PEG4 Dye

To prepare cells for subcellular localization studies, place coverslips in a Petri dish and add a culture medium to cultivate the cells. Allow the cells to grow until they reach the desired confluency. Then, carefully remove the old medium and introduce a staining solution that has been pre-warmed to 37°C. This staining solution should contain MitoTracker dye (1 µM), Cy5-PEG4 dye (1 µM), and Hoechst 33342 dye (1 µg/mL). Incubate the cells with this mixture for 30 minutes to ensure thorough staining.

After incubation, gently remove the staining solution and replace it with fresh, pre-warmed culture medium. Then, the stained cells were examined using a confocal laser scanning fluorescence microscope to observe the subcellular localization of the Cy5-PEG4 dye.

Utilize Image J software to analyze microscopy images. This software facilitates the examination of images captured during microscopy. Focus on quantifying the overlap of fluorescence signals from the MitoTracker probe with the Cy5-PEG4 dye. This step is crucial as it reveals the intracellular compartments the Cy5-PEG4 dye associates with, providing insights into its subcellular localization.

### Animals

Adult Sprague Dawley (SD) rats were acquired from Charles River Laboratories (Wilmington, MA). The rats were housed under a consistent 12-hour light/12-hour dark cycle in a temperature- and humidity-regulated room. They had free access to food and water throughout the study. The procedures strictly complied with the guidelines outlined in the National Institutes of Health’s Guide for the Care and Use of Laboratory Animals. The research protocol received approval from the Institutional Animal Care and Use Committee at Michigan Technological University.

### Primary Neuronal Cell Culture Procedure

Neuronal cells were extracted from the brains of 1-day-pold SD pups using a papain dissociation system (Worthington Biochemical Corporation, Worthington, NJ, USA) following the manufacture’s instruction. The extracted cells were cultured in Neurobasal-A medium enhanced with a 2% B27 supplement and a 1% combination of penicillin-streptomycin antibiotics. The culturing was conducted on 35 mm dishes pre-coated with poly D-lysine. These dishes were kept in a 5% CO_2_ incubator at 37°C, and half of the culture medium in each dish was refreshed every three days to ensure optimal growth conditions. After 7 to 10 days of culturing, the cells were exposed to various concentrations of the RNYIAQVD peptide for 48 hours. Following the exposure to the peptide, the neuronal cultures underwent a dual-staining process. They were stained with Cy5-PEG4 dye at a concentration of 1 µM, which produces a red, fluorescent signal, and Hoechst stain at a concentration of 0.1 µg/mL, emitting blue fluorescence. The purpose of this dual staining was designed to exam the changes in mitochondrial morphology and dynamics in response to the RNYIAQVD peptide treatment.

### *In vivo* Administering Cy5-PEG4 via Intraperitoneal Injection

To investigate whether Cy5-PEG4 can cross the BBB in rats, an intraperitoneal (i.p.) injection process is employed. Cy5-PEG4 is prepared in sterile phosphate-buffered saline (PBS). The SD rats are then anesthetized with a 5% isoflurane mixture, with careful monitoring to ensure a depth of anesthesia that minimizes distress. Once anesthetized, the rats are placed on a sanitized surface to administer the Cy5-PEG4 solution, delivered at a concentration of 200 nM (10 µl) through a sterile syringe and needle. 24 hours after the Cy5-PEG4 injection, the rats are euthanized with an overdose of isoflurane. This is followed by transcardial perfusion with PBS, which clears the blood vessels to prepare the brain for extraction. The harvested brain tissue is then fixed and sectioned for further examination.

The brain sections are analyzed under a confocal fluorescent microscope to determine the extent of Cy5-PEG4 penetration into the brain tissue. This experimental setup includes control groups that receive only the vehicle solution for comparison purposes. Using Image J software, the fluorescence intensity within various brain regions is quantified, enabling a thorough comparison and assessment of Cy5-PEG4’s ability to penetrate the BBB in the rats.

## Results

### Molecular Design Rationale

The molecular design of the fluorescent dye Cy5-PEG4 is a deliberate strategy to address the specific challenges of brain cell imaging, mainly targeting the mitochondria. The choice to use asymmetric cyanine as the core structure of Cy5-PEG4 is based on its superior photophysical properties, such as increased brightness and sensitivity, and better absorption and emission characteristics compared to symmetric cyanine dyes. Asymmetric cyanine dyes, with their longer wavelengths, provide the added advantage of deep tissue penetration and reduced background autofluorescence, crucial for brain imaging where clarity and depth are imperative [6]. A notable feature of asymmetric cyanine dyes is their large Stokes shifts, which minimize the overlap between excitation and emission light, thus improving signal clarity [7]. This property is pivotal in differentiating the dye’s signals from background noise and autofluorescence, which is especially beneficial in the complex environment of brain tissue. The integration of a PEG linker in the Cy5-PEG4 dye serves several purposes. Firstly, PEGylation enhances the water solubility and biocompatibility of the compound [8], crucial for an imaging agent that needs to circulate within the bloodstream and penetrate the brain tissue. This enhancement in pharmacokinetics is essential to ensure that the dye can reach the mitochondria within brain cells efficiently. The addition of PEG extends the systemic circulation time of the imaging agents, reducing renal clearance thus improving the chances of the dye to cross the BBB [9]. Furthermore, the terminal alkyne group introduces a bioorthogonal handle to the dye, enabling selective and efficient conjugation with biomolecules through click chemistry reactions, particularly Cu-AAC, as described by Wang et al. [10]. This aspect of molecular design facilitates the attachment of specific ligands or targeting moieties that can direct the imaging agents to mitochondrial targets within the brain cells. Cy5-PEG4 is designed to navigate the BBB — a challenging barrier due to its selective permeability. The small size of the terminal alkyne group on the Cy5-PEG4 reduces steric hindrance, allowing for better passage through the BBB without compromising the dye’s functionality once inside the brain. The near-infrared (NIR) emission property of the Cy5 component in Cy5-PEG4 is another critical aspect tailored for brain imaging. NIR emission enables acquiring the images from the deeper brain tissue with less autofluorescence interference [11]. This feature is essential for *in vivo* imaging, offering a window to observe mitochondrial dynamics within the brain cells. Cy5-PEG4’s molecular structure, featuring an asymmetric cyanine core for optimal photophysical properties, a PEG linker for improved biocompatibility and pharmacokinetics, and a terminal alkyne for bioconjugation and BBB penetration, is engineered to address the unique obstacles of brain imaging. Each component of Cy5-PEG4’s design contributes to its ability to reach and image mitochondria within brain cells effectively, illustrating the potential of this dye to advance the field of neurological imaging.

### Synthesis of the fluorescent Cy5-PEG4 dye

**Scheme 1.**
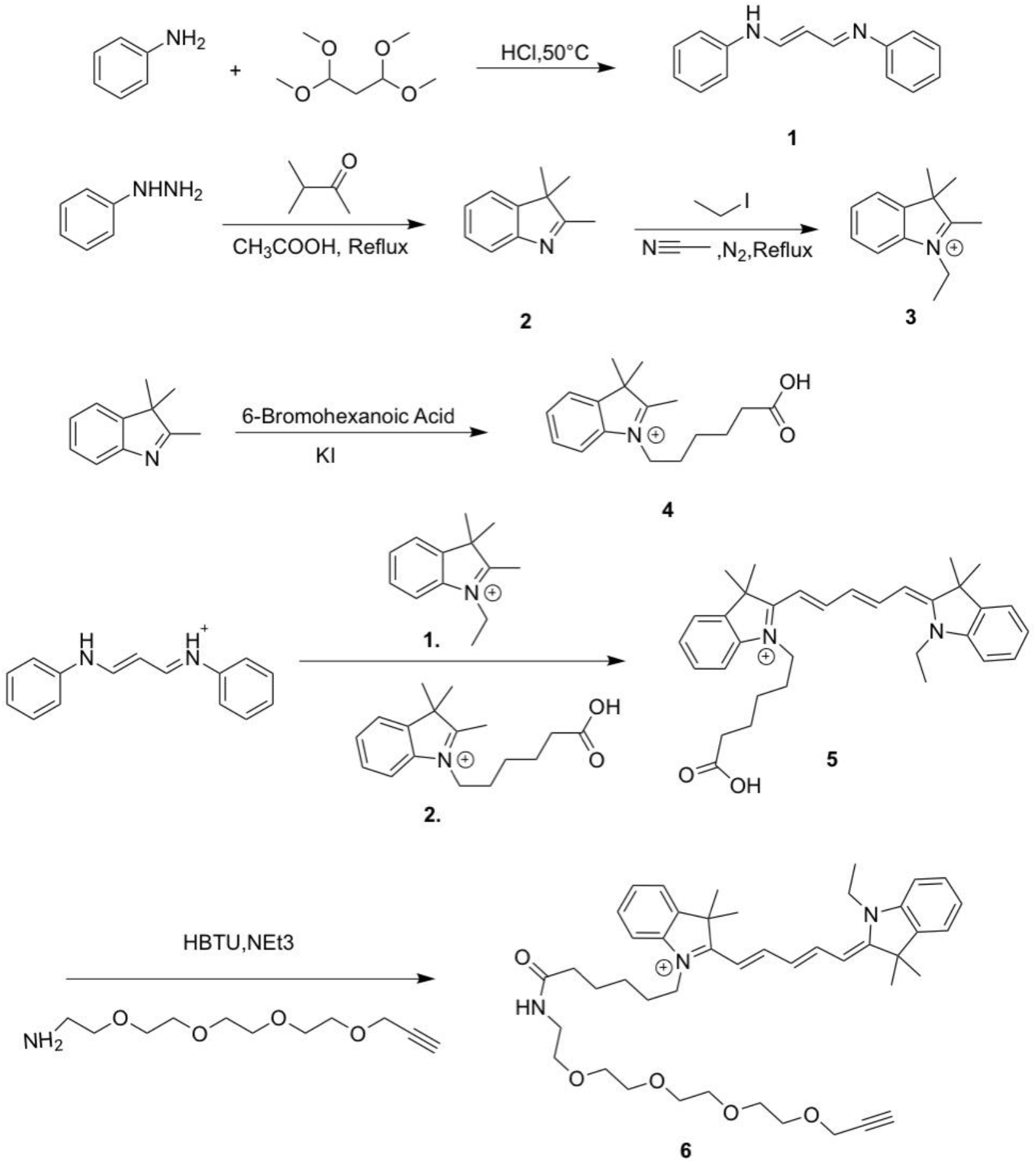
Synthesis of the fluorescent dye, Cy5-PEG4.

In pursuit of enhanced NIR fluorophores suitable for *in vivo* applications, a concerted synthetic approach was taken to develop the Cy5-PEG4 conjugate. The synthesis integrated a PEG linker to augment the conjugate’s solubility and biocompatibility. The synthetic route was initiated with the reaction of 1,1,3,3-tetramethoxypropane (TMP) and aniline in the presence of hydrochloric acid, producing compound **1**. Mechanistically, an iminium ion was generated via protonation, which underwent intramolecular cyclization. This was a critical step, requiring careful pH control to ensure the desired product’s formation. Base-mediated deprotonation led to rearrangement, forming a cyclized structure. Special attention was needed to prevent over-deprotonation, which could lead to undesirable by-products. The ensuing synthesis of compound **2** presented a condensation challenge. In an acidic environment, phenylhydrazine reacted with 3-methylbutanone to form an imine. Ensuring anhydrous conditions was crucial to avoid hydrolysis of the sensitive imine. This imine was protonated and rearranged into a more stable enamine that cyclized intramolecularly. Each step required precise temperature control to favor the correct pathway and avoid competing reactions. The formation of compound **3** entailed an SN2 reaction, utilizing the deprotonated nitrogen in compound **2** as a nucleophile. This species attacked ethyl iodide, necessitating a rigorously anhydrous acetonitrile medium to avoid competing hydrolysis of the alkylating agent. Careful monitoring of the reaction environment ensured the successful displacement of the iodide ion and the stabilization of the resultant charge on the nitrogen atom. Next, compound **4** was synthesized by nucleophilic substitution with 6-bromohexanoic acid in presence of KI. This intermediate rearranged to a zwitterionic form before losing a proton to yield the product. This step demanded stringent control of stoichiometry and reaction temperature to drive the formation of the zwitterionic species and avoid its decomposition. Constructing the asymmetric cyanine core of dye **5** was achieved through a condensation reaction, merging the carboxy-indoleninium salt **4** with compound **3** and malonaldehyde dianil. This process, which is vital for integrating the quaternary ammonium group, was delicate and required an inert atmosphere to prevent oxidation of the sensitive intermediates. Following this, an alkylation step was performed to functionalize the tertiary amino group, necessitating precise pH and temperature control to avoid over-alkylation. The culmination of this synthesis was the conjugation with a PEG moiety, which enhanced the dye’s hydrophilic profile and modulated its pharmacokinetics. This step had to be carried out under conditions that would prevent premature PEGylation and ensure a high-yield coupling to compound **5** to provide compound **6**, Cy5-PEG4.

### Spectroscopic Analysis of Cy5-PEG4 Dye

Cy5-PEG4 dye exhibits excellent absorption and emission properties with distinct, narrowly defined peaks, making it highly specific and effective for use in complex biological samples. The dye shows a notable Stokes shift, where the absorbance peak occurs at a much shorter wavelength than the emission peak, enhancing its ability to separate emission from excitation light, which is crucial for fluorescence applications (Figure 1).

**Figure 1:**
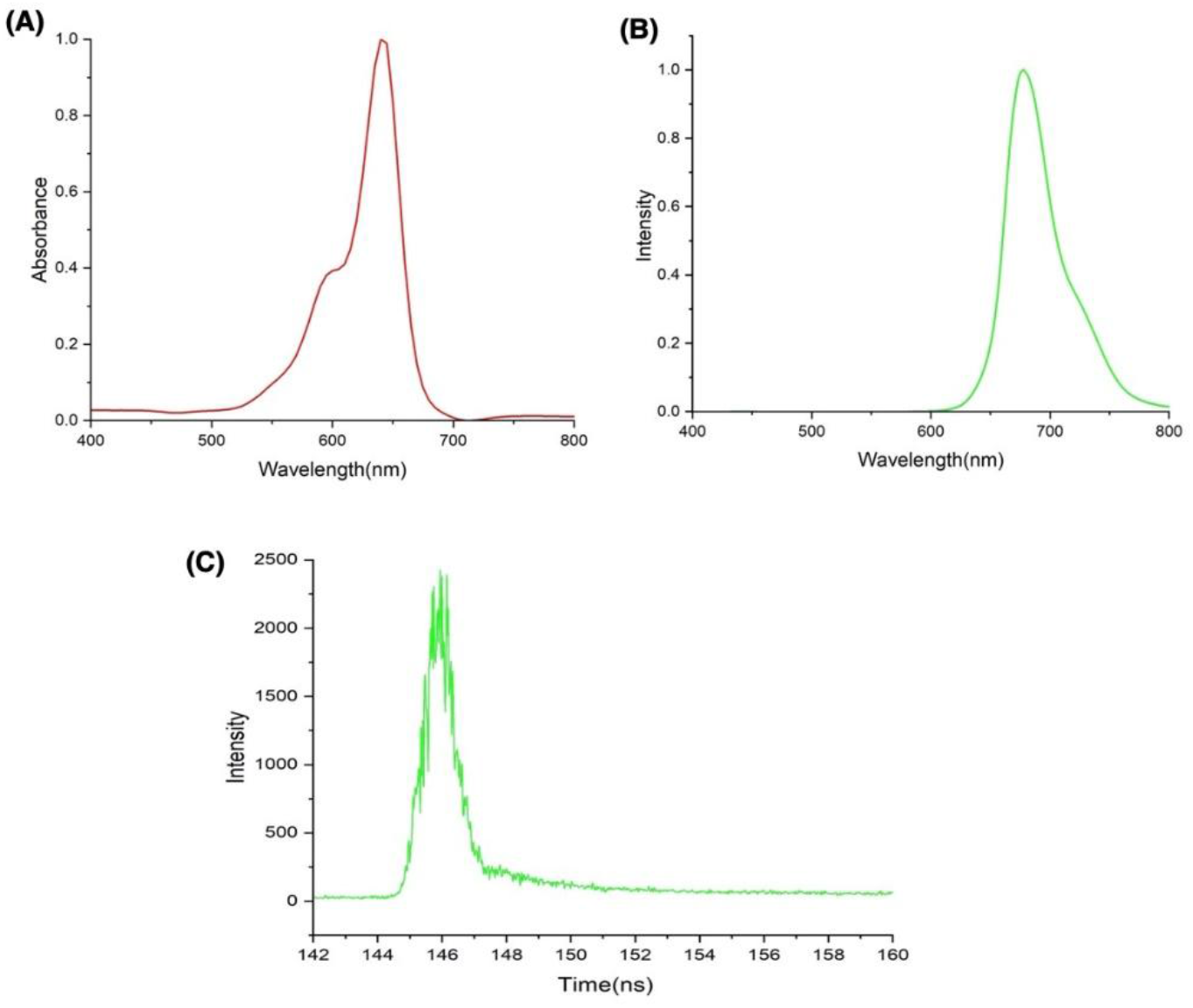
Cy5-PEG4 dye exhibits strong absorbance (**A**) and emission (**B**) in the red region, suitable for fluorescence applications like imaging or sensing. The short fluorescence lifetime (**C**) enhances its utility in time-resolved applications, making it a promising candidate for advanced optical techniques in biological systems.

The high intensity of the emission peak indicates a high fluorescence quantum yield, providing robust and detectable signals suitable for analytical and diagnostic methods. The sharp and symmetric peaks reflect the dye’s high purity, ensuring reliable experimental results. Positioned in the red to near-infrared spectrum, Cy5-PEG4 benefits from deeper tissue penetration and reduced scattering, ideal for *in vivo* imaging. The incorporation of PEG improves solubility and minimizes nonspecific binding, enhancing its performance in biological assays.

The fluorescence lifetime profile reveals consistent photophysical behavior, advantageous for imaging applications where uniform signal intensity is critical. The rapid fluorescence kinetics support applications like flow cytometry, where quick, precise analysis is required. Additionally, the distinct peaks and fast decay characteristics are valuable for time-resolved measurements, allowing for the isolation of Cy5-PEG4’s specific signal from background autofluorescence. Cy5-PEG4’s stability and resistance to photobleaching make it particularly useful in long-term imaging studies, maintaining a high signal-to-noise ratio and reducing background interference. This combination of properties ensures Cy5-PEG4’s effectiveness across various imaging techniques, including microscopy, flow cytometry, and *in vivo* studies, offering clear, consistent, and high-resolution results.

### Mitochondrial targeting property of Cy5-PEG4 dye

Our investigation of Cy5-PEG4’s mitochondrial localization employed HUVEC, which are widely used in vascular biology research. By mimicking BBB dynamics and angiogenesis, HUVEC extend their utility beyond cardiovascular studies, providing valuable insights into neurodegenerative disease mechanisms [12]. Confocal microscopy showed significant mitochondrial accumulation of Cy5-PEG4 in HUVEC, highlighting the dye’s potential for precise mitochondrial imaging. Similarly, HEK293 cells, derived from human embryonic kidney tissue, are frequently used in neurodegenerative research due to their neuronal-like characteristics [13]. These cells express over 60 neuronal genes, including neurofilament proteins, neuroreceptors, and ion channel subunits [14], making them suitable for studying neurological disorders. Consistent mitochondrial localization of Cy5-PEG4 was observed across HUVEC, HEK293, and HeLa cell lines (Figure 2). Quantitative analysis using ImageJ and Pearson’s correlation coefficient (PCC) [15] confirmed Cy5-PEG4’s strong mitochondrial colocalization, with a PCC of 0.88 in HUVEC, demonstrating high specificity for mitochondria. Similar results were observed in HEK293 and HeLa cells, with PCC values of 0.84 and 0.92, respectively, confirming the dye’s consistent mitochondrial targeting across various cell types. These results underscore the effectiveness of Cy5-PEG4 in targeted mitochondrial imaging and validate PCC as a reliable measure for assessing the precision of this fluorescent probe, supporting its use in diverse biological studies.

**Figure 2.**
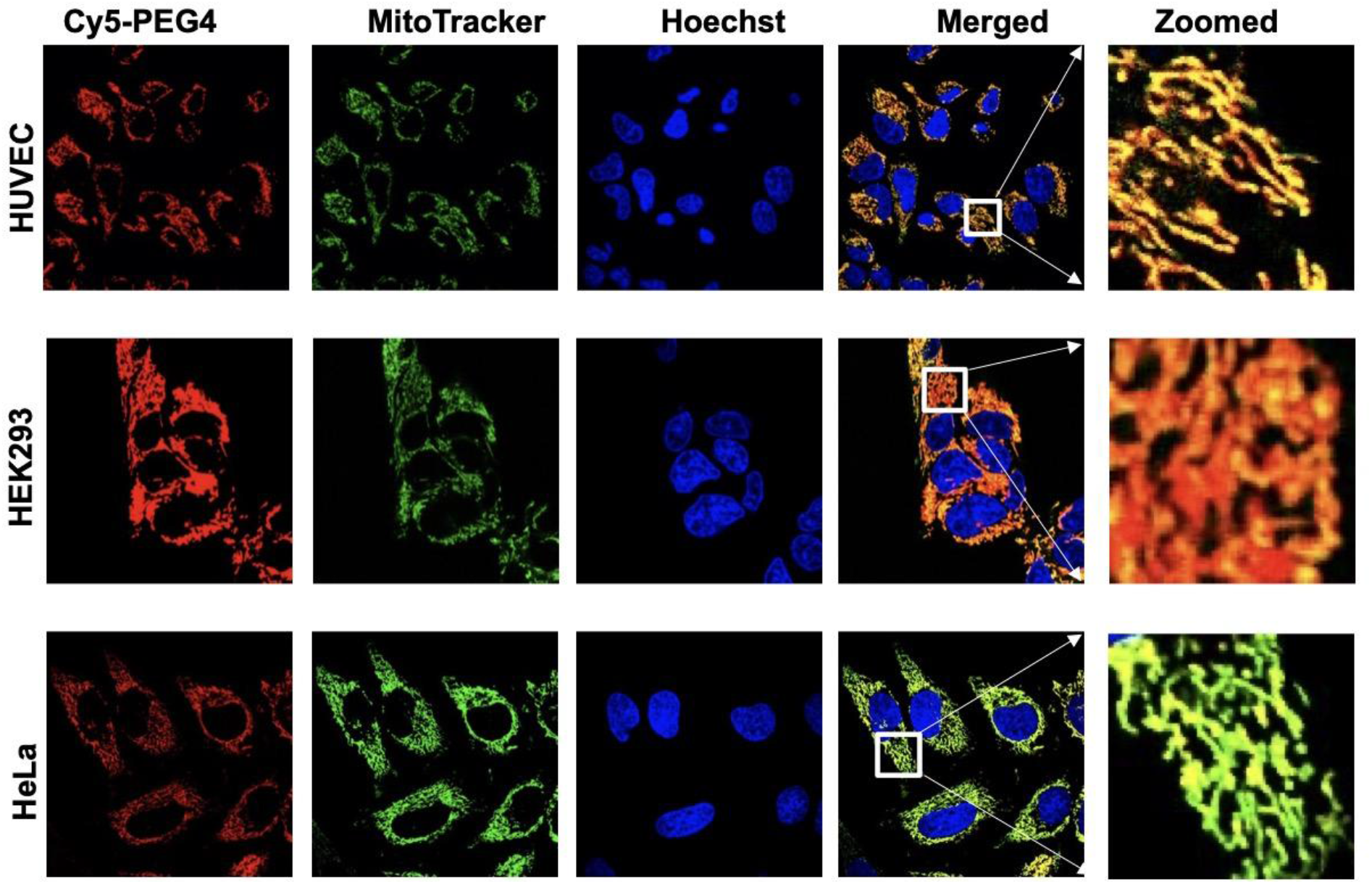
Intracellular distribution of the Cy5-PEG4 dye compared to MitoTracker. Three different cell lines were incubated with the Cy5-PEG4 dye (1 µM, red fluorescence), followed by counterstaining with MitoTracker (1 µM, green fluorescence), Hoechst 33342 (1 µg/mL, blue fluorescence) for 40 min; the images were taken at randomly selected areas to ensure the imaging conditions were consistent throughout the experiments. The experiments were repeated four times to ensure the reliability of the results. Cells were imaged on an inverted laser scanning fluorescent microscope (Olympus) using a 60 × oil immersion objective lens.

### Cy5-PEG4 can effectively image mitochondria in 3D cultured cells

Three-dimensional (3D) models bridge the gap between *in vitro* and *in vivo* systems by enhancing cellular differentiation and complexity, while also reducing the need for animal experiments [16]. In these models, cells can develop more complex structures, such as neurite outgrowths and cell aggregates, which closely resemble the architecture of neuronal tissues. This resemblance to *in vivo* conditions provides researchers with more physiologically relevant models for exploring various aspects of neurobiology and neuropharmacology.

In our study, we observed that MitoTracker was ineffective in labeling mitochondria in 3D-cultured PC12 cells (images not shown), highlighting several challenges associated with using this dye in 3D culture systems. MitoTracker’s limited penetration depth is a significant obstacle, particularly in dense 3D cultures or thick tissue samples. While MitoTracker efficiently labels mitochondria in 2D cultures, its performance declines as the depth of the sample increases, leading to incomplete labeling and poor visualization of mitochondria located deeper within the structure. Additionally, photobleaching poses another limitation, as the prolonged light exposure required for imaging in 3D cultures accelerates signal loss, reducing image quality over time.

Moreover, MitoTracker’s pH sensitivity complicates mitochondrial imaging in 3D environments. Although it targets mitochondria effectively at the acidic pH found within these organelles, variations in pH gradients within 3D cultures can influence labeling efficiency, compromising the accuracy of the data. Given these challenges, we explored Cy5-PEG4 as a potential alternative to improve mitochondrial imaging in 3D cultures.

Our findings showed that Cy5-PEG4 effectively labeled mitochondria in 3D cultured PC12 cells, demonstrating its unique advantages over MitoTracker (Figure 3). While MitoTracker’s variable efficacy across different cell models suggests that environmental factors in 3D cultures may impede its uptake, Cy5-PEG4’s enhanced solubility and biocompatibility, likely due to its PEG linker, allow it to penetrate cellular barriers and selectively label mitochondria in these complex environments. This makes Cy5-PEG4 a promising alternative, especially in conditions where conventional probes fall short, broadening the toolkit available for mitochondrial imaging.

**Figure 3.**
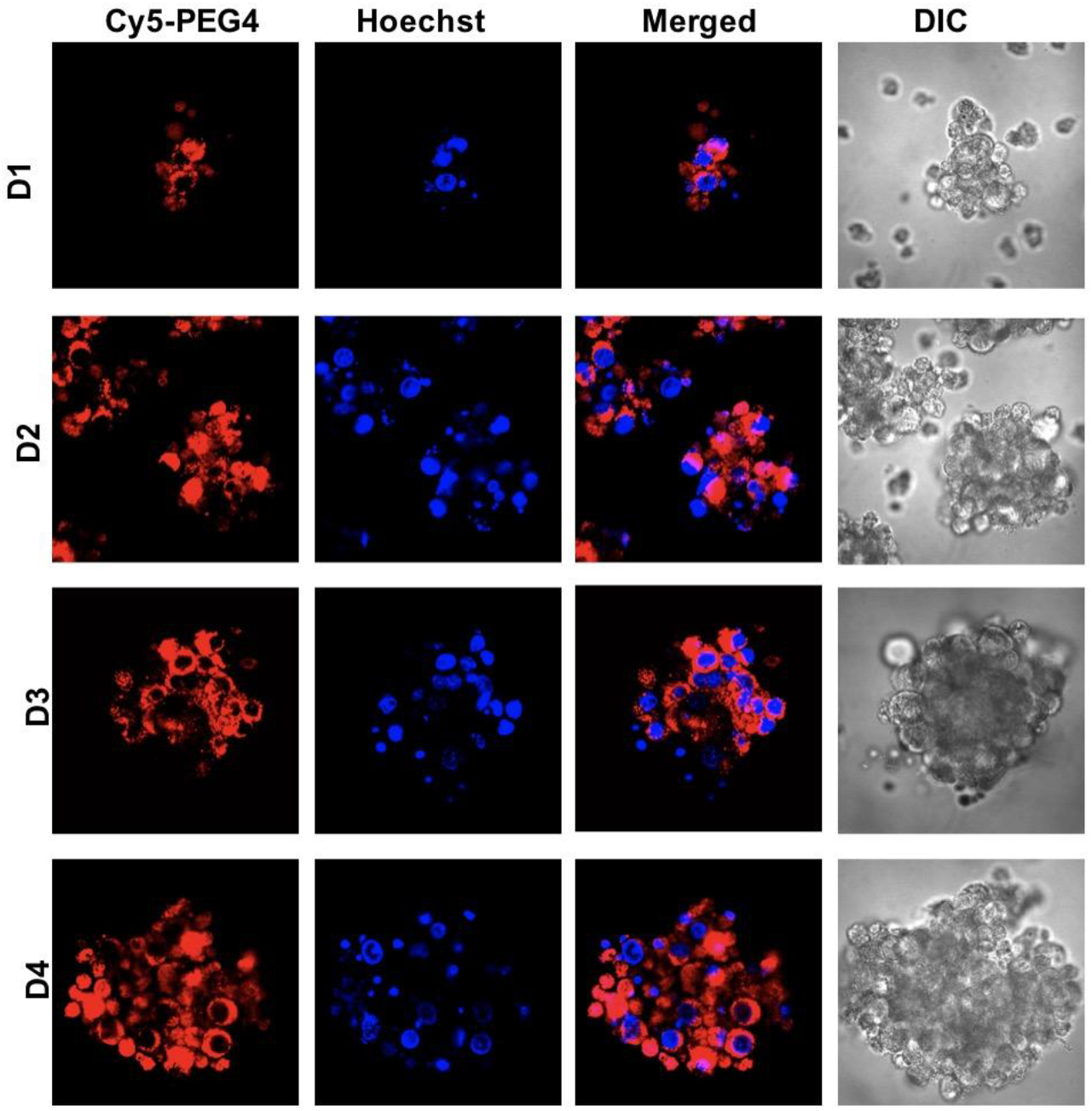
Mitochondrial imaging in 3D cultured PC12 cells using Cy5-PEG4 from day 1 (D1) to day 4 (D4) reveals dynamic changes in distribution, morphology, and functionality as the cell structures evolve. As the structures mature, imaging complexity increases, emphasizing the need for advanced techniques to accurately assess mitochondrial function and morphology throughout the 3D culture.

During the four-day 3D culture period, the cellular structures undergo distinct morphological transformations (Figure 3). Initially, cells seeded into systems such as low-attachment plates or hydrogels are mostly in suspension, forming small, irregular clusters with minimal extracellular matrix production. By the second day, intensified cell-cell interactions result in the formation of small spheroids or early organoid structures, with compacted cells establishing tight junctions and early signs of differentiation and hypoxia within the core.

As the cultures expand on the third day, the spheroids grow, with outer layers actively dividing and the inner core showing reduced proliferation due to limited nutrient and oxygen access. Extracellular matrix production increases, and tissue-specific markers begin to appear, indicating the development of proliferative and quiescent zones within the spheroids. By the fourth day, these structures become more complex, with clearly defined layers; the core often experiences hypoxia or necrosis, and the spheroids exhibit functional characteristics relevant to their intended models, such as hormone secretion and drug responsiveness.

Mitochondrial imaging throughout these four days reveals progressive changes in distribution, morphology, and function. On the first day, mitochondria are scattered and primarily elongated or fragmented within loosely organized clusters, allowing relatively straightforward imaging due to the low cell density. By the second day, mitochondria form more interconnected networks, particularly near active cell junctions, but increased cell density begins to introduce imaging challenges, such as light scattering and obstructions.

By the third day, mitochondrial organization reflects the developing cellular zones within the spheroids. Mitochondria in the outer layers remain active and networked, while those in the hypoxic inner core exhibit altered morphology, appearing rounded or fragmented. Imaging becomes more difficult due to light scattering and limited dye penetration. By the fourth day, the mature structures reveal marked differences between the outer, active regions and the dysfunctional core, further complicating imaging due to pronounced scattering and background noise.

The contrast between Mitotracker and Cy5-PEG4 highlights the importance of selecting mitochondrial probes that align with the specific dynamics of 3D culture environments. Cy5-PEG4’s ability to effectively image mitochondria across these stages demonstrates its potential as a robust tool for investigating mitochondrial dynamics, function, and morphology in complex cellular models, making it a valuable alternative when conventional probes do not perform optimally.

### The Cy5-PEG4 dye can effectively detect mitochondrial Morphology Alterations

COVID-19, primarily a respiratory disease, has also been associated with neurological symptoms such as memory loss, sensory confusion, severe headaches, and stroke, affecting up to 30% of cases and sometimes persisting as long COVID. These symptoms are believed to result from the virus infecting the central nervous system (CNS), though the exact molecular mechanisms remain unclear. Reynolds’ team identified two peptides from the SARS-CoV-2 proteome that form amyloid assemblies, which are highly toxic to neuronal cells, suggesting that cytotoxic aggregates of these viral proteins may contribute to the observed neurological symptoms [5].

The motivation behind using the novel mitochondrial-targeting fluorescent dye Cy5-PEG4 to study the effects of the RNYIAQVD peptide on primary neuronal cells stems from the urgent need to understand the neurological impact of COVID-19 [5]. Persistent neurological symptoms reminiscent of neurodegenerative diseases have been observed in COVID-19 patients [17], raising concerns that amyloidogenic peptides from SARS-CoV-2, which form toxic aggregates, could be contributing to these complications. Cy5-PEG4 enables a precise investigation into whether such peptides play a role in the neuropathology of acute COVID-19 and its prolonged aftermath, known as ‘long COVID.’

The RNYIAQVD peptide, derived from the SARS-CoV-2 spike protein, may significantly affect mitochondrial morphology and function during viral infection [5]. While direct studies on this peptide’s impact on mitochondria are limited, there are plausible mechanisms through which SARS-CoV-2 peptides could influence mitochondrial dynamics, given the virus’s broader effects on cellular function. Viral infections often lead to mitochondrial fragmentation due to increased reactive oxygen species (ROS) and inflammatory cytokines [18, 19], both elevated in SARS-CoV-2 infections. The RNYIAQVD peptide could contribute to this stress by interacting with mitochondrial proteins or modulating cellular signaling pathways. Disruption of the balance between mitochondrial fusion and fission, essential for healthy mitochondrial networks, may also occur, leading to altered mitochondrial morphology and cellular dysfunction.

Historical parallels from other amyloid-producing viruses support this investigation, as SARS-CoV-2’s ability to invade the nervous system, similar to predecessors like Zika and SARS-CoV-1, suggests that its peptides, like RNYIAQVD, might exert harmful effects on the brain. This study aims to reveal the specific interactions of the RNYIAQVD peptide with primary neuronal cells, focusing on mechanisms that could lead to neuronal injury.

The overarching goal is to clarify the influence of the RNYIAQVD peptide on mitochondrial integrity and functionality, which are crucial for neurocellular health. Cy5-PEG4 allows real-time observation of mitochondrial responses to the peptide, providing insights into potential viral triggers of neurodegeneration. This knowledge is expected to contribute significantly to developing therapeutic approaches to alleviate the neurological impacts of COVID-19, advancing efforts to address the persistent cognitive and neurological challenges faced by those recovering from the virus infection.

As shown in **Figure 4**, we observed that the RNYIAQVD peptide has a dose-dependent impact on mitochondrial morphology and dynamics in primary neuronal cells. At high concentrations (2 µM), the peptide leads to mitochondrial fragmentation, a characteristic observed in neurodegenerative diseases, possibly indicating a mechanism by which SARS-CoV-2 contributes to COVID-19-associated neurological symptoms. This fragmentation could signal cellular stress or damage, potentially playing a role in developing or worsening neurodegenerative processes associated with the long-term effects of COVID-19. At lower concentrations (0.05 µM), however, the peptide induced an adaptive response, promoting a more interconnected and tubular mitochondrial network, implying a protective cellular mechanism to counteract stress and maintain health.

**Figure 4.**
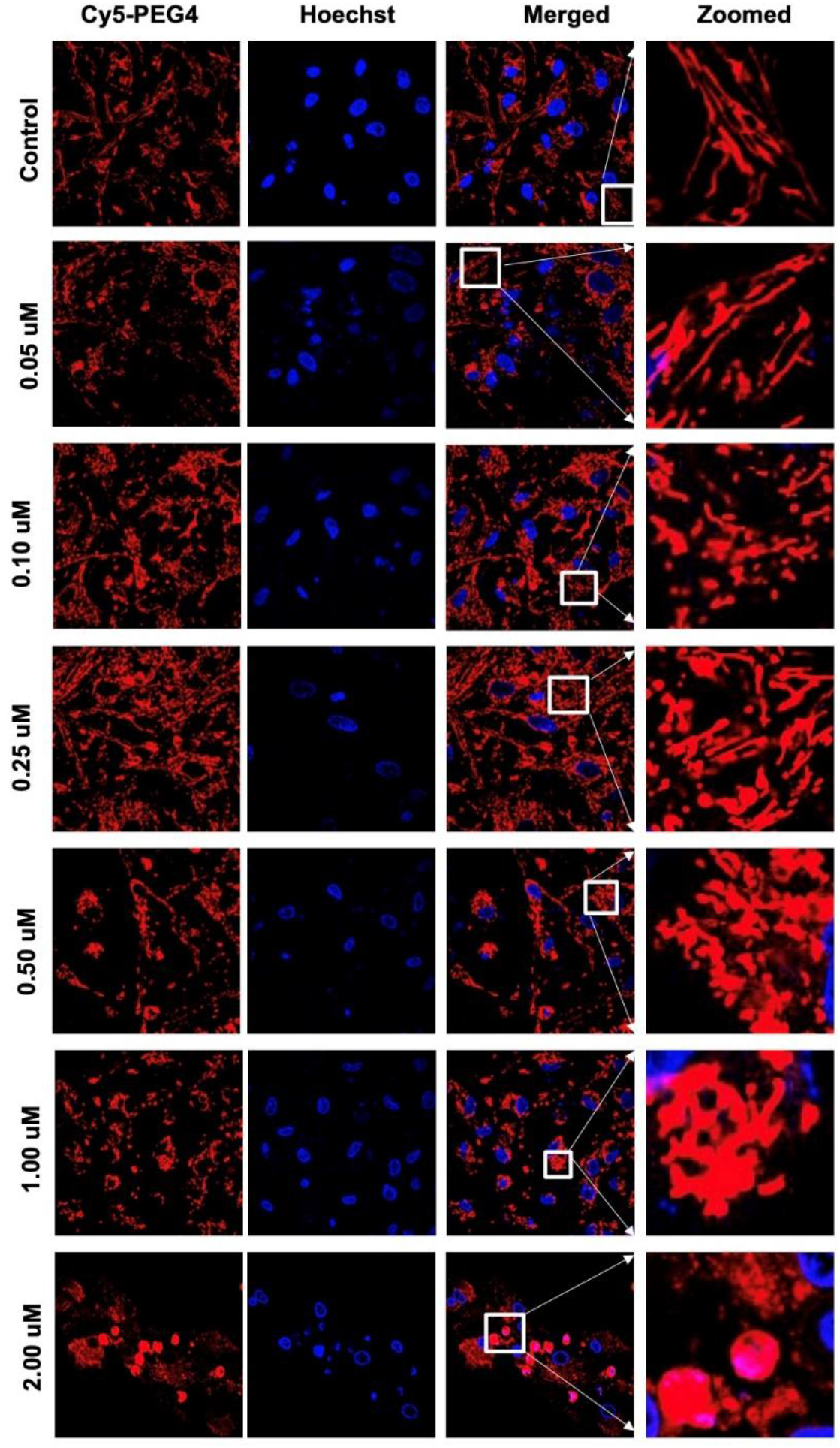
Representative confocal images of the primary neuronal cell lines in the presence of the RNYIAQVD peptide at varying concentrations (0, 0.05, 0.10, 0.25, 0.50, 1.00 and 2.00 µM) for 48 h. Cells were stained with Cy5-PEG4 (1 µM, red fluorescence) and co-stained with Hoechst (1 µg/mL, blue fluorescence). The images were taken at randomly selected areas to ensure the imaging conditions were consistent throughout the experiments. The experiments were repeated four times to ensure the reliability of the results. The fluorescent images were obtained with a confocal laser scanning fluorescent microscope using a 60 × objective lens.

The use of Cy5-PEG4 proved effective in detecting these mitochondrial changes, making it a valuable tool for monitoring mitochondrial alterations in real-time. This ability to visualize mitochondrial behavior could be instrumental in further understanding the pathophysiological effects of SARS-CoV-2 on neuronal cells and developing strategies to address the neurological aspects of COVID-19. Insights from this study may also shed light on the connection between viral infections and neurodegenerative disease processes.

## The Cy5-PEG4 dye can effectively traverse the BBB and image brain cells’ mitochondria

Our investigation into Cy5-PEG4’s BBB permeability and neurotoxicity focused on evaluating its effects on glial fibrillary acidic protein (GFAP) and ionized calcium-binding adapter molecule 1 (Iba1) expression, which are key markers of astrocyte and microglial activation, respectively. By monitoring these markers, we aimed to assess the dye’s biodistribution, pharmacokinetics, and biological interactions within the CNS, providing critical insights into its safety profile and biocompatibility.

To evaluate Cy5-PEG4’s impact within the CNS, we examined its ability to cross the BBB and its potential toxicological effects by analyzing GFAP and Iba1 expression following intraperitoneal administration. GFAP, a marker of astrocyte activation, increases in response to neuronal injury or inflammation, indicating astrogliosis and glial reactivity, processes commonly observed in neurodegenerative diseases [20, 21]. Similarly, Iba1 serves as a marker of microglial activation, reflecting the microglia’s response to neuronal damage and neuroinflammation [21, 22]. Our results showed no significant changes in GFAP and Iba1 levels in S.D. rats after Cy5-PEG4 administration, suggesting minimal neurotoxicity and a negligible inflammatory response, thereby supporting the safety of Cy5-PEG4 as a CNS imaging agent without triggering adverse glial activation or neuroinflammation (Figures 5, 6).

**Figure 5.**
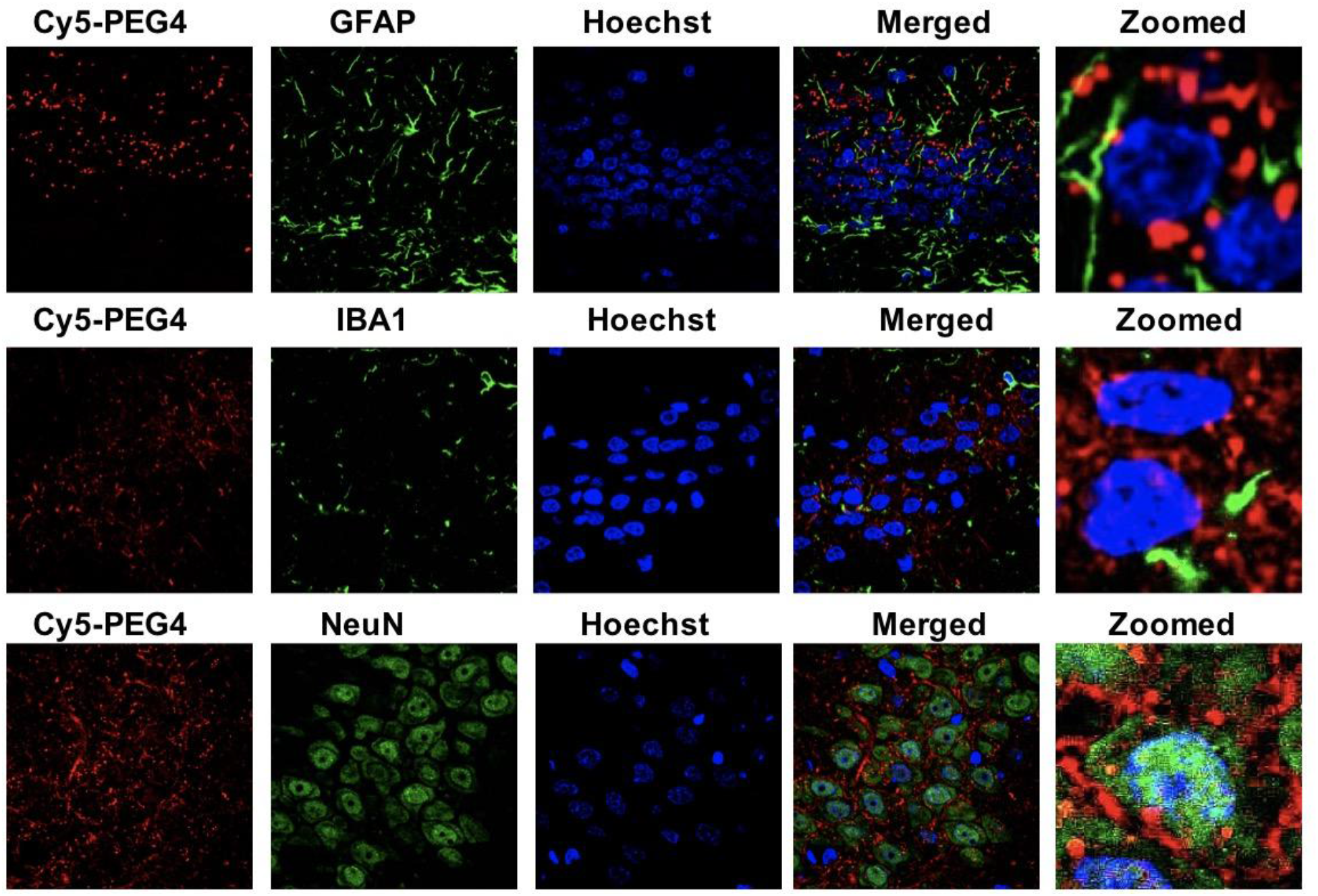
Visualization of the Cy5-PEG4 dye in the hippocampus regions after i.p. injection of Cy5-PEG4 (10 uM): (**A**) representative confocal laser scanning images of rat brain sections fixed with 4% PFA. For the entire figure, the red channels represent the fluorescent dye Cy5-PEG4; the green channel represents GFAP-(row I), IBA1-(row II), and NeuN-positive (row III) cells in the brain sections. Zoomed-in pictures are shown on the right. The fluorescent images were obtained with a confocal laser scanning fluorescent microscope using a 60 × objective lens. (**B**) Quantitative analysis of the fluorescence intensity of GFAP. (**C**) Quantitative analysis of the fluorescence intensity of IBA1.

**Figure 6.**
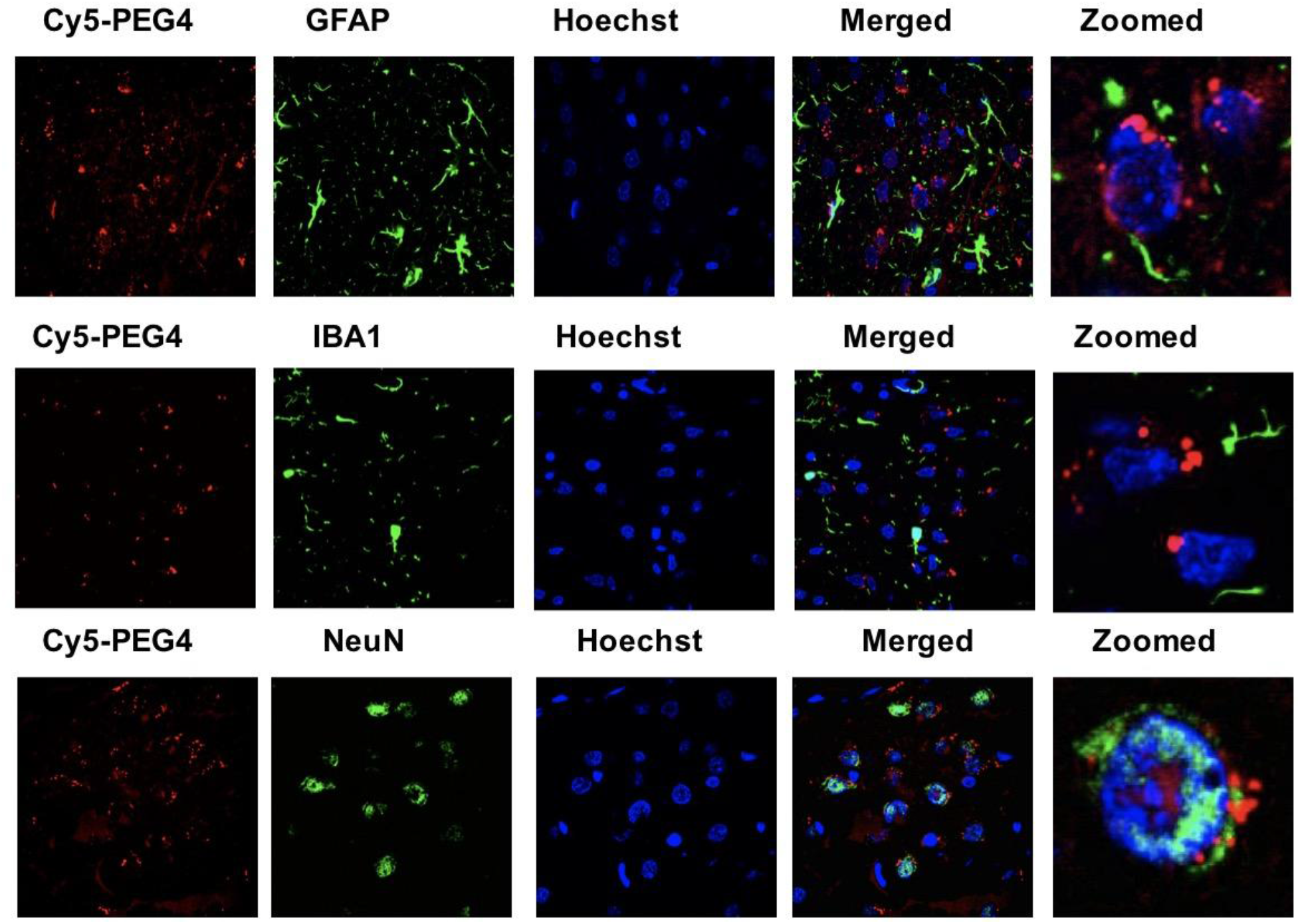
Visualization of the Cy5-PEG4 dye in the cortex regions after i.p. injection of Cy5-PEG4 (10 uM): (**A**) representative confocal laser scanning images of rat brain sections fixed with 4% PFA. For the entire figure, the red channels represent the fluorescent dye Cy5-PEG4; the green channel represents GFAP-(row I), IBA1-(row II), and NeuN-positive (row III) cells in the brain sections. Zoomed-in pictures are shown on the right. The fluorescent images were obtained with a confocal laser scanning fluorescent microscope using a 60 × objective lens. (**B**) Quantitative analysis of the fluorescence intensity of GFAP. (**C**) Quantitative analysis of the fluorescence intensity of IBA1.

Further analysis revealed that Cy5-PEG4 successfully crosses the BBB and selectively accumulates in NeuN-positive neurons, with minimal association with Iba1-positive microglia and GFAP-positive astrocytes. This selective neuronal localization highlights Cy5-PEG4’s potential as a specific neuronal imaging agent, offering valuable insights into neuronal behavior, neurodegeneration, and targeted pathologies. The specificity for NeuN, a neuronal marker, emphasizes Cy5-PEG4’s utility in neuroscience research, allowing precise imaging of neurons without significant interference from glial cells.

The limited interaction of Cy5-PEG4 with glial cells reduces the likelihood of confounding glial activation, enhancing its suitability for studies centered on neuronal physiology and pathology. This selectivity enables focused investigations of neuronal dynamics, such as metabolism, viability, and function, without the complications associated with glial reactivity. The ability to specifically label neurons positions Cy5-PEG4 as an ideal tool for exploring neuronal health and disease.

Cy5-PEG4’s BBB penetration and selective targeting of neurons underscore its potential for research into neurodegenerative diseases, providing insights into neuron-specific processes without inducing inflammatory or immune responses from glial cells. This property not only enhances its safety but also supports its use in non-invasive, repeated imaging or therapeutic applications in neurological studies.

The ability of Cy5-PEG4 to cross the BBB and selectively target neurons also suggests potential for therapeutic applications, such as neuron-specific drug delivery systems for treating neurological disorders that require targeted interventions. Overall, Cy5-PEG4’s unique properties make it a powerful tool for studying neuronal function and dysfunction, with broad applications in diagnostics, therapeutics, and advancing our understanding of the CNS’s neuronal environment.

### Novelty Statement

The development of Cy5-PEG4 as a mitochondrial-targeting fluorescent dye represents a significant advancement over the previously reported Cy5-PEG2 [9], addressing key limitations that restricted the latter’s use in complex biological systems, such as the brain. Cy5-PEG4 was specifically designed with a longer PEG linker to enhance its pharmacokinetic properties (data will be reported elsewhere), improving blood-brain barrier penetration and enabling superior mitochondrial targeting within brain tissue. These modifications allow for more precise, high-resolution imaging of mitochondrial dynamics in vivo, crucial for advancing research in neurodegenerative diseases and other neurological conditions. Additionally, Cy5-PEG4’s extended PEG linker reduces phototoxicity and improves stability in biological environments, making it particularly suitable for non-invasive, long-term imaging studies, including those requiring repeated sessions. This dye’s advanced physicochemical properties and superior safety profile—crossing the BBB without causing neuroinflammation or cytotoxicity—make it a safer and more versatile alternative for complex models, including 3D cultures and *in vivo* applications.

Cy5-PEG4 also expands the scope of research applications by effectively labeling mitochondria in neuronal cells exposed to neurotoxic peptides, such as those associated with SARS-CoV-2, yielding novel insights into mitochondrial dysfunction in COVID-19-related neurodegeneration. Its high sensitivity and specificity in detecting mitochondrial changes make it an invaluable tool for early biomarker discovery and therapeutic evaluation in various pathological conditions. These unique attributes establish Cy5-PEG4 as a critical asset not just for imaging but also for advancing the understanding of mitochondrial alterations in disease progression, supporting the development of targeted therapeutic strategies.

## Conclusion

This study highlights the significance of Cy5-PEG4 as a novel mitochondrial-targeting dye that facilitates the exploration of neurological effects associated with COVID-19 and other neurodegenerative diseases. Cy5-PEG4’s ability to cross the BBB without inducing neuroinflammation or toxicity underscores its potential as a safe and effective tool for *in vivo* brain imaging, enabling detailed investigations of mitochondrial responses to neurotoxic peptides, including those from SARS-CoV-2. The dye’s capacity to capture dynamic changes in mitochondrial distribution and function in maturing 3D cultured cells further emphasizes the need for advanced imaging tools to accurately study these complex processes. Our findings support the broader application of Cy5-PEG4 in neurobiological research, diagnostics, and therapeutic development, demonstrating its critical role in advancing our understanding of mitochondrial dynamics in health and disease, and highlighting its relevance in addressing the neurological impacts of COVID-19 and other neurodegenerative conditions.

## Limitation

While this study highlights the promising role of Cy5-PEG4 in investigating the neurological effects of COVID-19, there are several limitations that should be acknowledged. First, the study primarily relies on *in vitro* models, including 3D cultured cells, which, while valuable, do not fully replicate the complexities of *in vivo* systems. This limitation underscores the need for further *in vivo* studies to validate the findings and assess Cy5-PEG4’s behavior within the living brain environment, including its long-term safety and biodistribution. Additionally, the interactions of Cy5-PEG4 with the RNYIAQVD peptide were studied in a controlled environment, which may not fully capture the variability of physiological conditions encountered in clinical scenarios. The specificity of Cy5-PEG4 for mitochondrial imaging under different pathological states, especially beyond the context of COVID-19, also remains to be thoroughly explored. Finally, while Cy5-PEG4 demonstrated minimal neuroinflammation or toxicity in initial assessments, comprehensive toxicological evaluations across diverse models and dosages are essential to establish its broader applicability and safety in neurological research and potential therapeutic use.

## ACKNOWLEDGMENT

We sincerely thank the NIH (R01HL163159, Z.S.), NIH (R15 EB035866, L.B), GLRC-ICC (L.B., R01805) and the American Heart Association (AHA) (grant 1807047, L.B.) for their generous financial support. We also extend our great gratitude to Dr. Rick Koubek for his encouragement and unwavering support throughout our project.

